# Sound source localization with varying amount of visual information in virtual reality

**DOI:** 10.1101/489484

**Authors:** Axel Ahrens, Kasper Duemose Lund, Marton Marschall, Torsten Dau

## Abstract

To achieve accurate spatial auditory perception, subjects typically require personal head-related transfer functions (HRTFs) and the freedom for head movements. Loudspeaker-based virtual sound environments allow for realism without individualized measurements. To study audio-visual perception in realistic environments, the combination of spatially tracked head mounted displays (HMDs), also known as virtual reality glasses, and virtual sound environments may be valuable. However, HMDs were recently shown to affect the subjects’ HRTFs and thus might influence sound localization performance. Furthermore, due to limitations of the reproduction of visual information on the HMD, audio-visual perception might be influenced. Here, a sound localization experiment was conducted both with and without an HMD and with a varying amount of visual information provided to the subjects. Furthermore, interaural time and level difference errors (ITDs and ILDs) as well as spectral perturbations induced by the HMD were analyzed and compared to the perceptual localization data. The results showed a reduction of the localization accuracy when the subjects were wearing an HMD and when they were blindfolded. The HMD-induced error in azimuth localization was found to be larger in the left than in the right hemisphere. Thus, the errors in ITD and ILD can only partly account for the perceptual differences. When visual information of the limited set of source locations was provided, the localization error induced by the HMD was found to be negligible. Presenting visual information of hand-location, room dimensions, source locations and pointing feedback on the HMD revealed similar effects as previously shown in real environments.

## Introduction

Virtual environments (VE) and virtual reality (VR) systems enable the study of audio-visual perception in the laboratory with a higher degree of immersion than obtained with typical laboratory-based setups. Head-mounted displays (HMDs) may allow the realistic simulation of visual environments, and loudspeaker-based virtual sound environments can reproduce realistic acoustic environments while maintaining the subjects’ own head-elated transfer functions (HRTFs). Combining HMDs and loudspeaker-based virtual sound environments could, therefore, be valuable for investigating perception in realistic scenarios.

To localize a sound source in the horizontal plane (azimuth) as well as in the vertical plane (elevation; see [1] for a review), three major cues are crucial: interaural time differences (ITDs), interaural level differences (ILDs), and monaural spectral cues generated by reflections from the body and the pinnae. An alteration of these cues can lead to a reduced localization accuracy [e.g. 2,3,4]. Wearing an HMD alters the sound localization cues [5, 6] and might reduce sound source localization accuracy.

Another factor that can affect people’s sound source localization ability is visual information. The ventriloquism effect describes the capture of an acoustic stimulus by a visual stimulus [7, 8], altering the perceived location of the acoustic stimulus. Maddox et al. [9] showed that the eye gaze modulates the localization accuracy of acoustic stimuli.

They found an enhanced ITD and ILD discrimination performance when the eye gaze was directed towards the source. Since HMDs have a reduced field-of-view relative to the human visual system, sound source localization abilities might also be affected due to reduced visual information. Furthermore, when having the room and the hand-location visible, the localization error has been shown to be lower than when subjects are blind-folded [10].

Modern proprietary VR systems have been shown to reproduce immersive visual environments and to provide reliable spatial tracking accuracy, both for the reproduction of virtual visual scenes as well as for headset- and controller-tracking [11, 12]. However, Niehorster et al. [12] showed that when the tracking system of the HMD is lost, for example due to the user blocking the path between the tracking system and the HMD with their hands, the VR system fails to maintain the correct spatial location. Consequently, the calibrated location of the HMD within the room can be offset, i.e. a certain direction in VR does not correspond to the corresponding direction in the real-world. Such offsets can be difficult to track in proprietary VR systems. To overcome this issue, a real-world to virtual-world calibration is proposed.

The aim of the present study was to clarify how sound source localization ability is affected by a VR system, and in particular, how HMD-induced changes of the binaural cues and virtual visual information alter sound localization accuracy. In order to address this, a sound localization experiment was conducted in a loudspeaker environment with and without an HMD. For the localization task, a hand-pointing method was employed utilizing commercially available hand-held VR controllers. To evaluate the accuracy of the pointing method, a visual localization experiment was conducted, where the subjects’ task was to point to specific locations. Since previous studies showed that sound localization accuracy can be influenced by visual information [10, 13], the amount of visual information provided to the subjects was varied in the VR environment. One condition included no visual information (blind-folded); another provided visual cues regarding the room dimensions and the hand-location; in a third condition, the subjects saw the loudspeaker locations and were additionally provided with a laser pointer for pointing on the perceived sound location. It was hypothesized that effects as observed in previous audio-visual localization experiments in real environments, may also be reflected in VR.

## Methods

### Subjects

Ten subjects (three female, seven male) with an average age of 24 years participated in the study. None of the subjects had seen the experimental room before. The subjects were blind-folded when they were guided into the room. They had normal audiometric thresholds equal to or below 20 dB hearing loss at the octave-band frequencies between 250 Hz and 8 kHz and self-reported normal or corrected vision. Nine of the ten subjects were right handed and were instructed to use their main hand to hold the controller. The data of the left-handed subject were mirrored on the median plane. For each subject, the interpupillary distance was measured and the HMD was adjusted accordingly to ensure a clear binocular image and to minimize eye strain.

### Acoustic reproduction method

The acoustic reproduction system consisted of 64 loudspeakers (KEF LS50, KEF, Maidstone, United Kingdom) housed in an anechoic chamber. The loudspeakers were arranged in a full sphere, at a distance of 2.4 m to the listening position, and driven by sonible d:24 (sonible GmbH, Graz, Austria) amplifiers. Twenty-seven of the loudspeakers in the frontal hemisphere were used in the present study. The loudspeakers were placed at three heights, at 0° (ear level) and ±28° elevation. Thirteen of the loudspeakers were at ear-level, distributed between −90° and +90° azimuth, with 15° separation. Seven loudspeakers were elevated by +28°, and seven by −28°, distributed between ±90° azimuth with 30° separation. All loudspeakers were equalized in level, delay and magnitude response as measured at the listening position. The loudspeakers were labelled using color coding (yellow, red and blue) to indicate elevation and numbers for azimuth locations (see Figure 1). The numbers ranged from one to thirteen, starting at −90° (left). The elevated loudspeakers at ±28° used only odd numbers, such that equal numerals were used for loudspeakers with the same azimuth angle.

**Fig 1.**
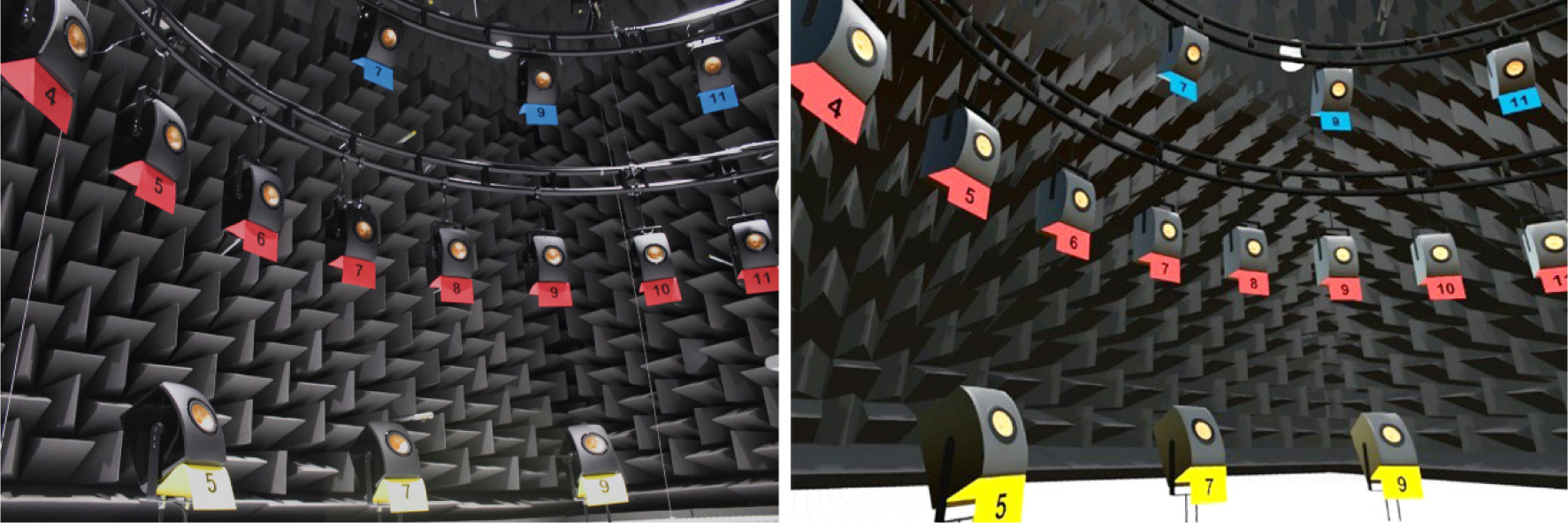
Photography (left) and screenshot (right) of the acoustic reproduction system in the real and in the virtual environment (RE and VE). The loudspeakers are numbered in azimuth and color coded in elevation.

### Visual reproduction method

The real environment (RE), shown in Figure 1 (left panel), was replicated to ensure comparable visual information with and without the HMD (Figure 1, right panel). To present the VE, the HTC Vive system (HTC Corporation, New Taipei City, Taiwan) was used. This system consists of an HMD and two handheld controllers to interact with the VE. Three additional Vive Trackers (HTC Corporation, New Taipei City, Taiwan) were used for VE-to-RE calibration (see details below). The spatial position and rotation of all devices were tracked with the infrared ray-tracking system. Blender (Blender Foundation, Amsterdam, The Netherlands) and Unity3D (Unity Technologies, San Francisco, CA) with the SteamVR plugin (Valve Corporation, Bellevue, WA) were used to replicate and present the VE, respectively.

When the aim is to replicate a real scenario in VR, while maintaining the interaction with real objects, it is crucial to ensure spatial alignment between the real and the virtual world. To calibrate the virtual world to the real world, the three Vive Trackers were positioned on top of the ear-level loudspeakers at 0° and ±45° azimuth. Discrepancies in the positions of the trackers in the RE and the VE were accounted for as follows:

1. The normal of the plane spanned by the three points corresponding to the positions of the trackers was calculated for the RE and for the VE and the difference in rotation between them was applied to the VE. This ensured the correct orientation of the VE.
2. To correctly position the VE, the difference between one tracker and the respective reference point in the VE was calculated and the VE was shifted accordingly. This resulted in an alignment of the RE and the VE at the chosen reference point.
3. The final rotation offset around the normal vector was corrected by calculating the angle difference of the vectors from the aligned reference point to an unaligned reference point in the VE and RE, i.e. to the known location of one of the other trackers and its virtual position.

After this procedure, the VE was aligned in both position and rotation relative to the RE. This method continuously accommodated for potential spatial discrepancies that might have occurred from tracking losses, as described by Niehorster et al. (2017). The system was recalibrated when either the tracker position error relative to the true position exceeded 2 cm or when the HMD lost tracking. The maximum allowed positional offset of the reference points resulted in a worst-case rotation error of 3.2°.

### Acoustic stimuli

The acoustic stimulus consisted of a pink noise burst with a duration of 240 ms and 20 ms tapered cosine ramps at the onset and offset. The noise burst was created in MATLAB (The Mathworks, Natick, MA) using a sampling frequency of 48 kHz. For each stimulus presentation, a new noise burst was created. The stimulus was presented at a sound pressure level (SPL) of 65 dB, and roved by values between ±3 dB, drawn from a uniform distribution. The short duration limits effect of head movements during the stimulus presentation [14] and the roving minimizes a spatial cue provided by directional loudness differences [15, 16]. The subjects were asked to re-center their viewing direction before each stimulus representation, i.e., to face the 0° on-axis loudspeaker at ear level. The HMD rotation was logged in a subset of the conditions and for a subset of the subjects to evaluate if, on average, the viewing direction was centered at the time of the acoustic stimulus exposure. An initial azimuth rotation of the HMD of −1°±3° standard deviation was found.

### Experimental conditions

Table 1 shows an overview of the eight experimental conditions considered in this study. The column ‘visual information’ shows the visual environment that was presented to the subjects. The stimulus refers to the localization task, which was either visual localization (visual search) or sound localization. The last column indicates whether the HMD was worn or not. Each condition and each of the 27 source locations (see section Acoustic reproduction method) was presented five times to each of the subjects. Thus, each condition consisted of 135 stimuli which were presented in fully random order.

**Table 1.** Overview over the conditions considered and grouped into blocks. Conditions were randomized within blocks. The conditions varied in available visual information, target stimulus and if the head-mounted display was worn.

| Block | Visual information | Stimulus | HMD |
| --- | --- | --- | --- |
| I | Blind-folded | Acoustic | No |
|  | Blind-folded | Acoustic | Yes |
| II | Virtual env., no loudspeaker (LS) | Acoustic | Yes |
| III | Virtual env., LS | Visual | Yes |
|  | Real env. | Visual | No |
|  | Real env. | Acoustic | No |
|  | Virtual env., LS | Acoustic | Yes |
| IV | Virtual env., LS, laser pointer | Acoustic | Yes |

The four blocks shown in Table 1 were presented in a fixed-order from Block I to Block IV to control for the exposure to the the loudspeaker locations. The conditions were randomized within the blocks. No pre-experimental training was conducted to avoid any bias of the subjects with respect to a specific condition or regarding visual information.

The two blind-folded conditions in Block I were used to examine whether, or to what extent, simply wearing the HMD has an influence on sound source localization. The reference localization accuracy was measured using the acoustic stimuli described above, while subjects were blind-folded with a sleeping mask. To assess the effect of the HMD frame on sound source localization, subjects wore the HMD on top of the sleeping mask.

The visual localization conditions in Block III were employed to investigate the baseline accuracy of the pointing method using the hand-held VR controller in the RE and the VE (Figure 1). The subjects were instructed to point at a loudspeaker location shown either on an iPad Air 2 (Apple Inc., Cupertino, CA) in the real environment or on a simulated screen within the VE.

To investigate the influence of the HMD on sound localization when visual information regarding the loudspeaker positions is available, the sound localization accuracy was evaluated in the real and in the virtual loudspeaker environments (Block III). The subjects were informed that sounds could also come from positions in-between loudspeakers. While the visual information regarding the loudspeaker positions was available in both conditions, the VE provided reduced visual information. The field-of-view of the HTC Vive is about 110° while the visual field of the human visual system is larger. Furthermore, in the VE only the hand-held controller was simulated but not the arm, which was inherently visible in the RE.

In addition to evaluating the pointing accuracy and the HMD-induced localization error, the effect of varying the amount of visual information on sound localization in the VE was investigated. In Block II, the experimental room (anechoic chamber) without the loudspeakers was rendered on the HMD and the subjects were asked to localize the acoustic stimuli as described for the previous conditions. The experiment thus included conditions with various degrees of visual information available to the subjects in the VR: no visual information, a depiction of the empty room including hand-location, and a depiction of the complete room including the locations of the sound sources.

To assess the role of visual feedback of the pointer location, in Block IV, a simulated laser pointer emerging from the hand-held controller was shown while the subjects completed the localization task in the VE with the room and the loudspeaker setup visible.

### Pointing method

The controller of the VR system was used as a pointing device. The subjects were instructed to indicate the perceived stimulus location by stretching their arm with the wrist straight while holding the controller, in an attempt to minimize intra- and inter-subject variability in pointing. The pointing direction was defined by the intersection point of an invisible virtual ray originating at the tip of the controller extending the base of the device and an invisible virtual sphere with the origin at the listeners head position and the same radius as the loudspeakers (r=2.4m). The perceived position of the source was indicated with a button press using the index finger.

On each trigger button press, the PC running the VR system transmitted an Open Sound Control (OSC) message via User Datagram Protocol (UDP) over an Ethernet network to the audio processing PC. The audio processing PC subsequently presented the next acoustic or visual stimulus, with a delay of three seconds to allow the subject to re-center the viewing direction. With a responding OSC message, the audio processing PC permitted the reporting of the perceived location after the playback completed.

A virtual model of the controller was rendered in all conditions containing visual information in the HMD. Thus, the visual feedback of the controller position in Blocks II and III was similar, independent of whether the HMD was worn or not. To standardize the pointing method for all audio-visual conditions, a direction marker, functioning as a visual pointing aid, was not provided in this study, except in the last condition (Block IV, Table 1), since a sufficiently comparable method was unfeasible in the real environment. Thus, the pointing method in Blocks II and III was similar to free-hand pointing.

### Physical analysis

The effect of the HMD on the acoustic ear signals was analyzed from B&K Head and Torso Simulator (HATS; Type 4128-C; Brüel & Kjær A/S, Nærum, Denmark) measurements with and without the HTC Vive HMD. Binaural impulse responses were recorded from all 64 loudspeakers with a 22-s long exponential sweep [17] in a frequency range from 60 Hz to 24 kHz and truncated to 128 samples (2.7 ms) to remove reflections from other loudspeakers and objects in the room. The dataset of the measurements can be found in a repository (zenodo.org/record/1185335). Acoustic perturbations of the HMD on the frequency spectrum were analyzed for the same set of loudspeakers as employed in the perceptual experiment by calculating the power in auditory filters between 200 Hz and 16 kHz with equivalent rectangular bandwidths [18] using the Auditory Modeling Toolbox [19]. The power in the auditory filters was averaged in three frequency regions from 200 Hz to 1 kHz, 1 to 5 kHz and 5 to 16 kHz.

Spectral differences (SD) were calculated as the mean absolute power differences of the three frequency regions, measured with and without the HMD. Interaural level differences (ILD) were determined in the same frequency region as the SD using the power differences at the output of the auditory filters between the left- and the right-ear signals. Interaural time differences (ITD) were calculated as the delay between the left- and right-ear signals. The delay of each impulse response was defined as the peak of the cross-correlation between the impulse response and its minimum-phase version [20].

### Pointing bias

It was hypothesized that subjects might have a bias in pointing direction due to the shape of the hand-held controller, and because they had no knowledge on where the ‘invisible ray’ was emerging from the controller. A bias for each subject was therefore calculated in azimuth and elevation as the mean of all source locations in the two visual localization conditions (real and virtual). Individual responses were then corrected by the calculated azimuth and elevation biases for all conditions except the condition with visual feedback of the pointer location (laser pointer condition).

### Analysis of behavioral responses

Statistical analyses on the subject response errors were performed by fitting mixed-effects linear models to the azimuth and elevation errors. The subject responses were corrected by a bias estimation due to the pointing method, as described above. Responses that were localized farther than 45° from the target location in either azimuth or elevation were treated as outliers. Of the 10800 subject responses, 0.29% were treated as outliers and discarded from the analysis.

Only the sources in the horizontal plane were considered in the statistical analyses of the azimuth localization errors. The azimuth stimulus location and the experimental condition (see Table 1) as well as their interaction were treated as fixed effects. Regarding the elevation error, the stimulus location in both azimuth (only azimuth directions that occurred in all elevation directions) and elevation, the experimental condition, as well as their interactions were treated as fixed effects. The influence of the subjects, the repetitions and their interaction were considered as random effects. The p-values were calculated based on likelihood-ratio tests for the random effects and on F-tests for the fixed effects [21]. Post-hoc analyses of within factor comparisons were performed using estimated marginal means calculated from the mixed-effects linear models and using Bonferroni p-value adjustments [22].

## Results

### Pointing Bias

Figure 2 shows the pointing bias in azimuth (squares) and elevation (circles) for each subject calculated from the visual localization experiments. Regarding the azimuth bias, the subjects tended to point slightly too far to the left (−3.5° to −0.1°), except for subject S07, who had a slight positive (right) azimuthal bias of 1.3°. Overall, the average bias across subjects in azimuth was −1.6° (left). The only left-handed subject (S08) showed a similar bias as the other subjects. The bias in elevation angle (circle) was found to be higher than the azimuth bias for all subjects, with an average value of 19.0°. The subjects generally tended to point too high. The variance across subjects was between 12.8° and 28.6° and is likely to be related to the shape of the hand-held controller and the internal reference of each subject of where the “invisible ray” emerges from the controller. The responses of the subjects were corrected by the pointing bias for all conditions except the laser pointer condition.

**Fig 2.**
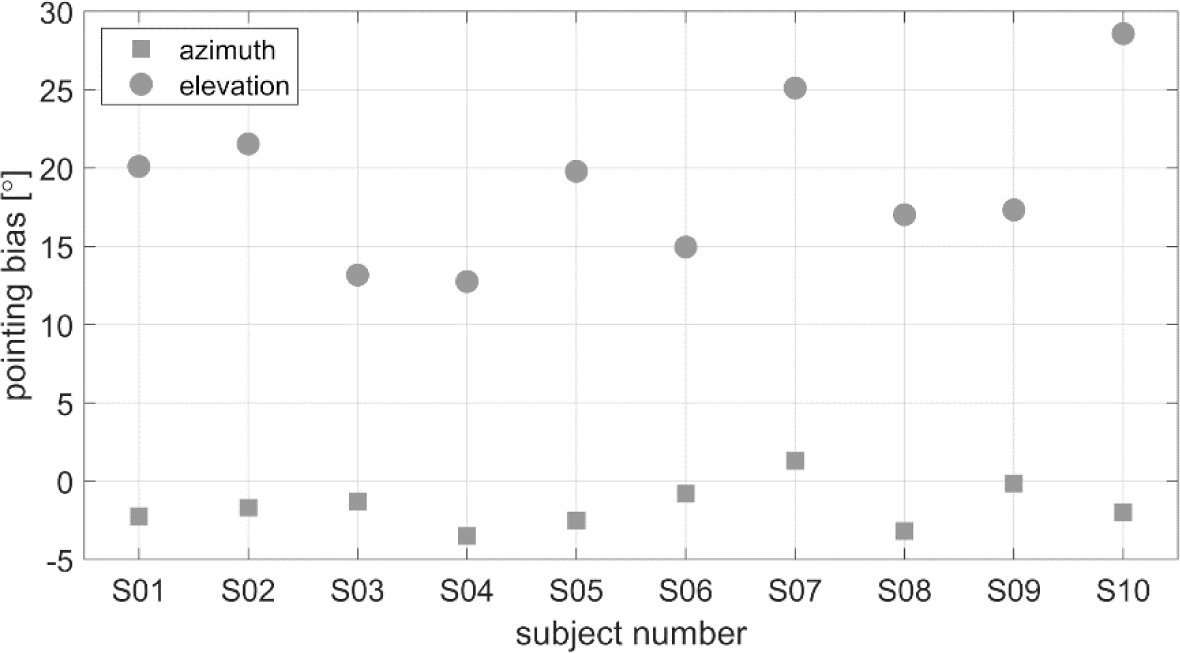
Pointing bias in azimuth (squares) and elevation (circles) for each subject. The bias was calculated as the mean error over all source locations in the two visual localization conditions. Negative angles indicate biases to the left and downwards for azimuth and elevation, respectively.

### Spectral differences and interaural errors

Figure 3 shows the spectral difference (SD) of the left-ear signals obtained with and without the HMD on the B&K HATS for the azimuth source locations between −90° and +90° and for the elevation angles of −28° (downward triangles), 0° (diamonds) and 28° (upward triangles). For ipsilateral sources (negative azimuth angles), the SD was low for all frequency regions and all elevation angles. In the low-frequency region between 200 Hz and 1 kHz (dashed lines), the SD was also below 1 dB for contralateral sources (positive azimuth angles), independent of the elevation angle. The SD in the mid-frequency (dashed-dotted lines) and high-frequency (solid lines) regions was found to be up to 6.3 dB for elevation angles at and above the horizontal plane (0°, 28°). For these elevations, the error was higher in the high-frequency region than in the mid frequency region, except at 60°. For mid- and high-frequency sources below the horizontal plane (i.e. an elevation angle of −28°), the SD was lower than for the other two elevation angles.

**Fig 3.**
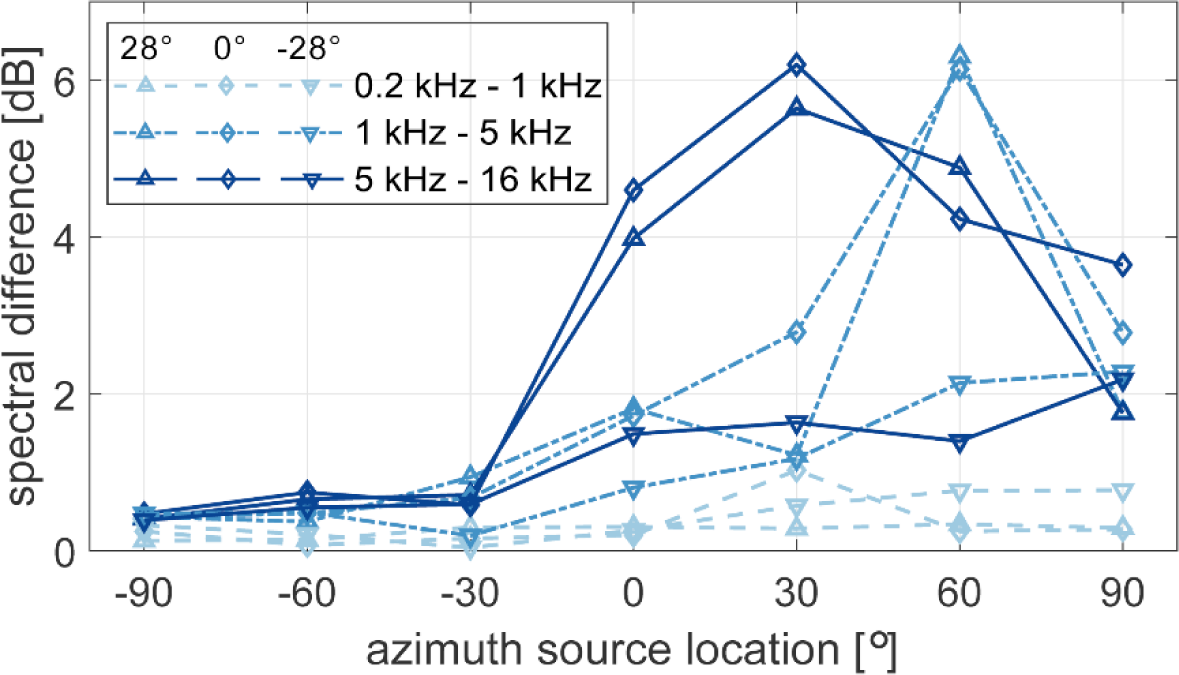
Spectral difference (SD) measured at the left ear of the B&K HATS with and without the HTC Vive. The angles in the legend represent the elevation angles considered in the current study. The SD was calculated in auditory bands and averaged over three frequency regions at low-, mid- and high frequencies as shown in the legend.

Figure 4 shows the signed errors in ILD (left panel) and ITD (right panel) induced by the HMD measured in the horizontal plane as a function of source azimuth angle. The ILD error in the low-frequency (dashed line) region was below 2 dB. In the mid- and high-frequency regions (dashed-dotted line and solid line), the error was lowest at the frontal source location (0° azimuth) and at 90. The largest error was about 6 dB at source angles of 60° and 30°, for the mid and high frequency regions, respectively. The ITD error was below 1 sample (21 µs) for source angles between 0° and 30° and increased to 62.5 µs for the source angle of 75°. The ITD error was 0 µs at 90° azimuth angle.

**Fig 4.**
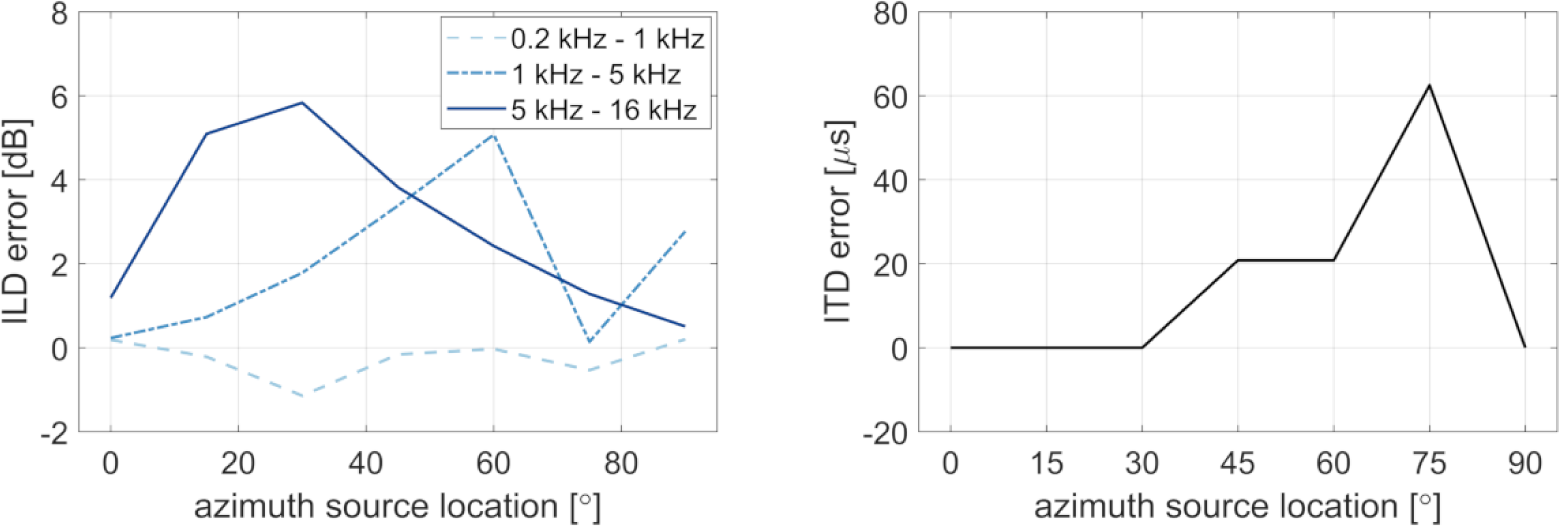
Signed errors of interaural differences in level and time (ILD and ITD) with respect to azimuth angles on the horizontal plane (0° elevation). Positive errors indicate larger ILDs and ITDs with than without the HMD. The ILDs were calculated in auditory bands and averaged over three frequency regions at low-, mid- and high frequencies as shown in the legend. The ITDs were calculated from the delays between the broadband binaural impulse responses (see Methods for details).

### Pointing accuracy with VR controllers

Figure 5 shows the responses of the subjects in the two visual localization experiments in the RE and the VE. In both conditions, the subjects’ task was to point to the center of the loudspeaker indicated on a screen. The filled black squares represent the 27 source locations. The small colored symbols represent the individual responses of the subjects and the large open black symbols indicate the mean responses, averaged across subjects and repetitions. The connecting lines between the target location and the mean responses indicate the localization error. The subjects generally pointed close to the correct loudspeaker, whereby the precision of the responses was generally higher for azimuth than for elevation localization.

**Fig 5.**
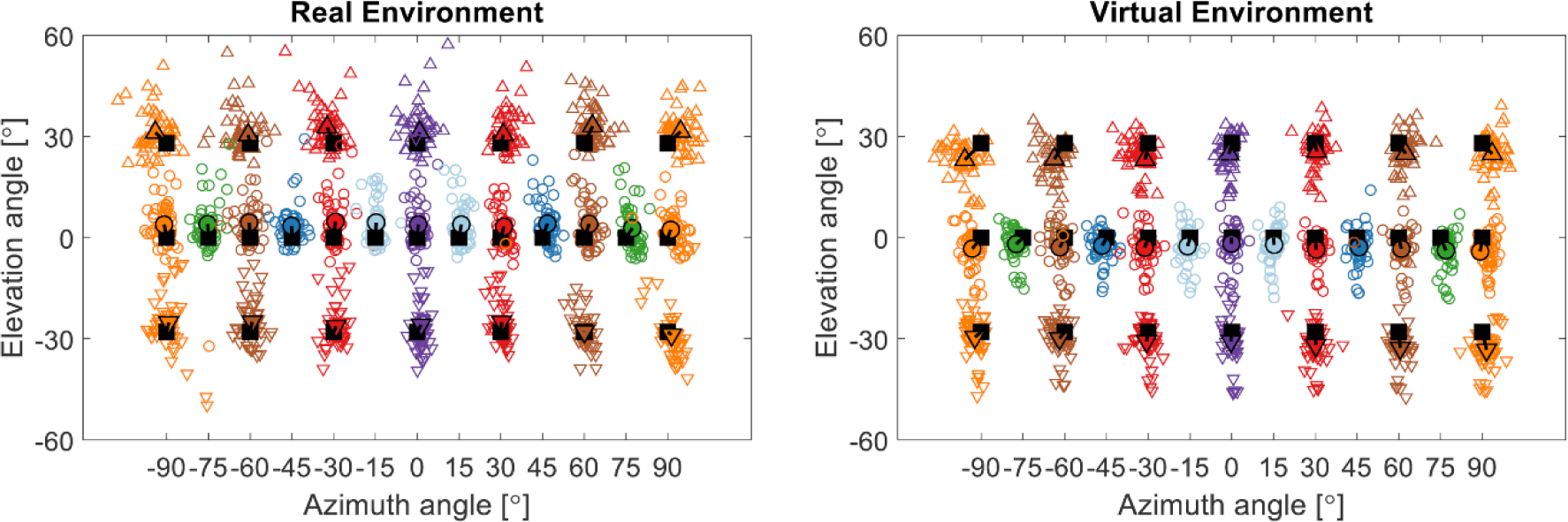
Response plot of the visual localization experiment for the real and virtual environment. The black squares represent the source locations. The small markers show the responses of the subjects and the large markers the mean response over subjects and repetitions. Negative angles represent sources to the left and downwards for azimuth and elevation, respectively.

Figure 6 shows the azimuth error obtained in the visual pointing experiment. The circles represent the mean absolute error and the boxplots indicate the signed error. The signed azimuth error was significantly affected by the conditions [F(1,1263)=60.63, p<0.0001], the azimuth source location [F(12,1263)=28.02, p<0.0001] and their interaction [F(12,1263)=1.9, p=0.0305]. The difference between the VE and the RE was only significant for sources on the left at −90° [t(1263)=4.52, p=0.0001], −75° [t(1263)=4.26, p=0.0003] and −30° [t(1263)=3.71, p=0.0028]. At −45° and −60° the difference was smaller and did not lead to significant effects after correction for multiple comparisons [-45°: t(1263)=2.64, p=0.1078; −60°: t(1263)= 2.73, p=0.0833]. Generally, the responses showed a small overshoot, i.e. the sources on the right side showed a shift to the right, while the responses for sources on the left tended to show a shift to the left. The overshoot was larger for sources on the left in the virtual environment with the HMD than in the real environment.

**Fig 6.**
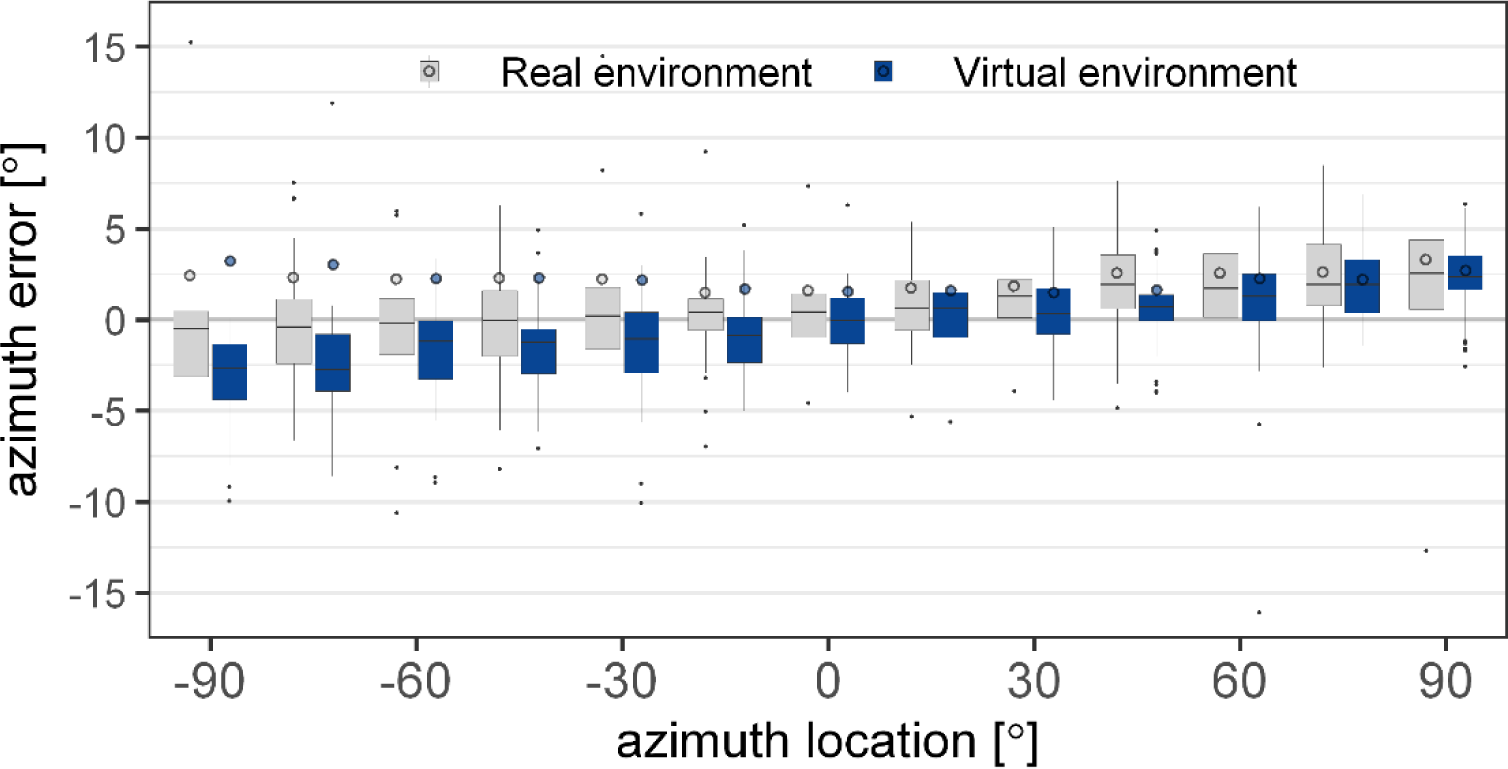
Mean absolute (circles) and signed (boxplots) azimuth error for visual localization in the virtual (dark blue) and the real (light grey) environment. The error is shown over the thirteen azimuth angles in the horizontal plane (0° elevation). The boxplots indicate the median (line) and the first and third quartile. The whiskers extend to 1.5 times the interquartile range.

The condition (RE vs. VE) was not found to have an effect on the absolute azimuth error [F(1,1263)=0.76, p=0.38]. Azimuth location [F(12,1263)=7.27, p<0.0001], as well as the interaction between the condition and azimuth location [F(12,1263)=1.83, p=0.0392], were found to be significant. However, after correction for multiple comparisons, no difference between the RE and the VE was found for any azimuth location [p > 0.11].

The analysis of the absolute elevation error showed that the conditions [F(1,2042)=9.77, p=0.0018] and the three-way interaction of condition, azimuth location and elevation location [F(12,1263)=2.02, p=0.0196] were significant. The two-way interaction of the conditions with the source locations in azimuth [F(6,2042)=2.21, p=0.0398] was significant, but not with the source locations in elevation [F(2,2042)=2.29, p=0.1]. The interaction between the source locations in azimuth and elevation was not significant [F(12,2042)=1.42, p=0.15] and also the source locations [Azimuth: F(6,2042)=0.99, p=0.43; Elevation: F(2,2042)=1.02, p=0.36] did not reveal significant effects.

The analysis of the signed elevation error showed significant contributions of the conditions [F(1,2069)=609.11, p<0.0001], the source azimuth [F(6,2069)=2.42, p=0.0245], the source elevation [F(2,2069)=7.85, p=0.0004] as well as the interactions between the source elevation and the conditions [F(2,2069)=9.51, p<0.0001] and both source locations [F(12,2069)=2.7, p=0.0013]. The interaction between the source azimuth and the conditions [F(6,2069)=0.75, p=0.61] and the three-way interaction [F(12,2069)=0.51, p=0.91] were not found to be significant. The difference between the RE and the VE was significant at all three source elevations, at 0° elevation [t(2069)=15.48, p<0.0001], as well as below [t(2069)=10.76, p<0.0001] and above [t(2069)=16.52, p<0.0001] the horizontal plane. The signed elevation error was positive (upwards) in the RE and negative (downwards) in the VE, as indicated by the lines between the target markers (black squares) and the response markers (colored squares) in Figure 5. On average, the subjects pointed 6.5° higher in the RE than in the VE.

### Influence of HMD on azimuth error

Figure 7 shows the absolute (circles) and the signed (boxplots) azimuth error as a function of the azimuth source locations. Negative angles represent sources on the left and positive angles indicate sources on the right of the subjects. The light grey boxes and circles represent results for the condition where the subjects were blind-folded and the dark blue boxes and circles show the results where the subjects were blind-folded and wore the HMD. The mean absolute azimuth error was always found to be larger with the HMD except for 0°. This difference was larger on the left than on the right side. The analysis of the signed error, employing a linear mixed effects model, showed that the effect of the conditions was not significant [F(1,1265)=0.6, p=0.44], while the source locations [F(12,1265)=62.04, p<0.0001] and the interaction [F(12,1265)=3.04, p=0.0003] were significant factors.

**Fig 7.**
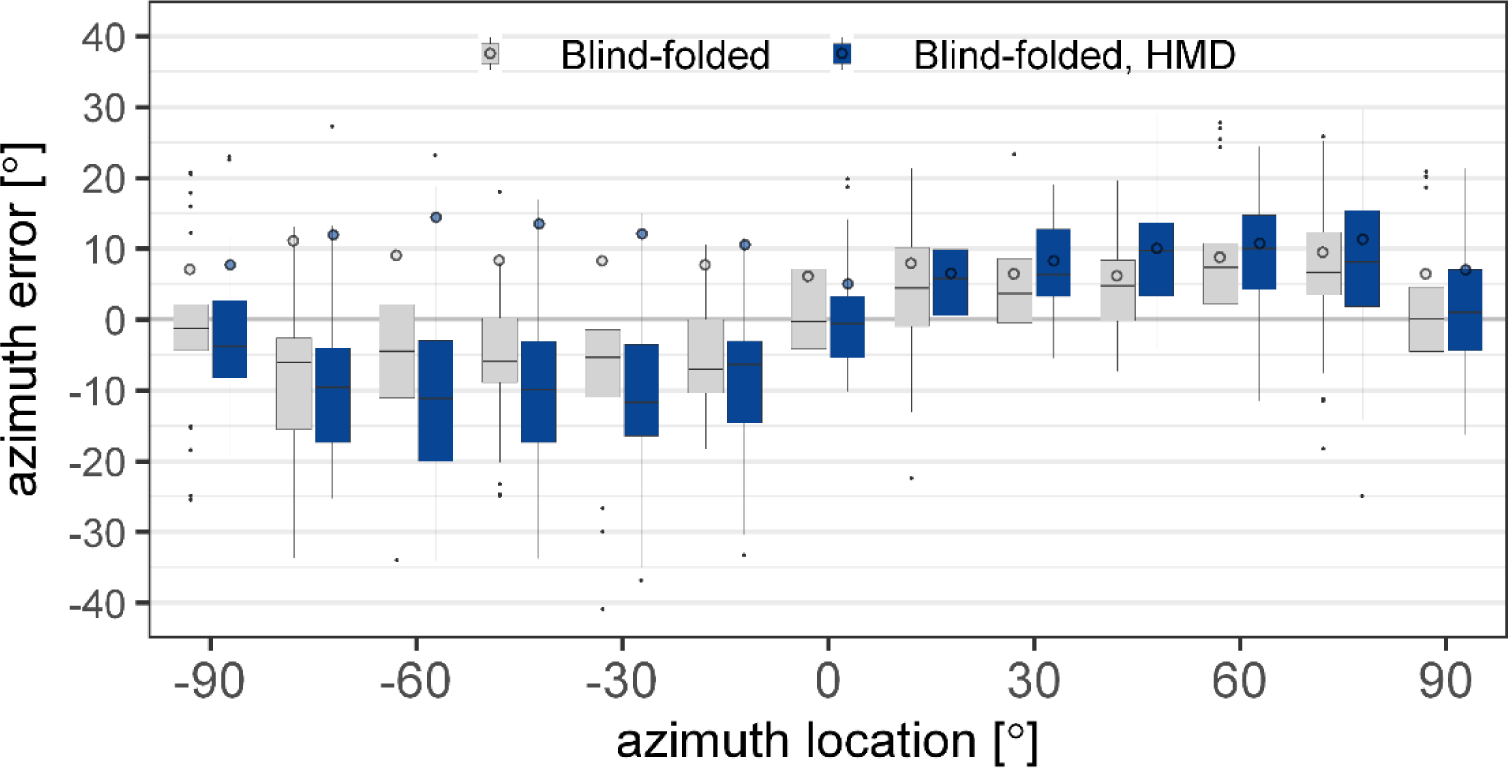
Mean absolute (circles) and signed (boxplots) azimuth error for acoustic localization of blind-folded subjects with (dark blue) and without (light grey) the head mounted display (HMD). The error is shown over the thirteen azimuth angles in the horizontal plane (0° elevation). The boxplots indicate the median (line) and the first and third quartile. The whiskers extend to 1.5 times the interquartile range.

The median of the signed error showed that the sources were perceived further lateral with the HMD than without. The post-hoc analysis for the sources on the left side showed that the difference between the conditions reached significance only at a source angle of −60° [t(1265)=3.3, p=0.0059]. At −45° the p-value exceeded the 5% significance level after correction for multiple comparisons [t(1265)=2.4, p=0.0944]. On the right side, the difference was significant only at 45° [t(1265)=-3.1, p=0.0119]. The error induced by the HMD on the signed azimuth error was larger on the left than on the right side at 45° [t(1265)=3.9, p=0.0014] and 60° [t(1265)=2.96, p=0.0374].

### Influence of visual information on azimuth error

Figure 8 shows the absolute (circles) and signed (boxplots) azimuth error as a function of the azimuth source locations for five conditions with varying visual information. The subjects were either blind-folded with the HMD (dark blue), were provided with information regarding the room dimensions and the hand position information (blue), could see the real room (light blue), were provided with the virtual version of the real room (white), or were provided with a laser pointer in the virtual room (grey).

**Fig 8.**
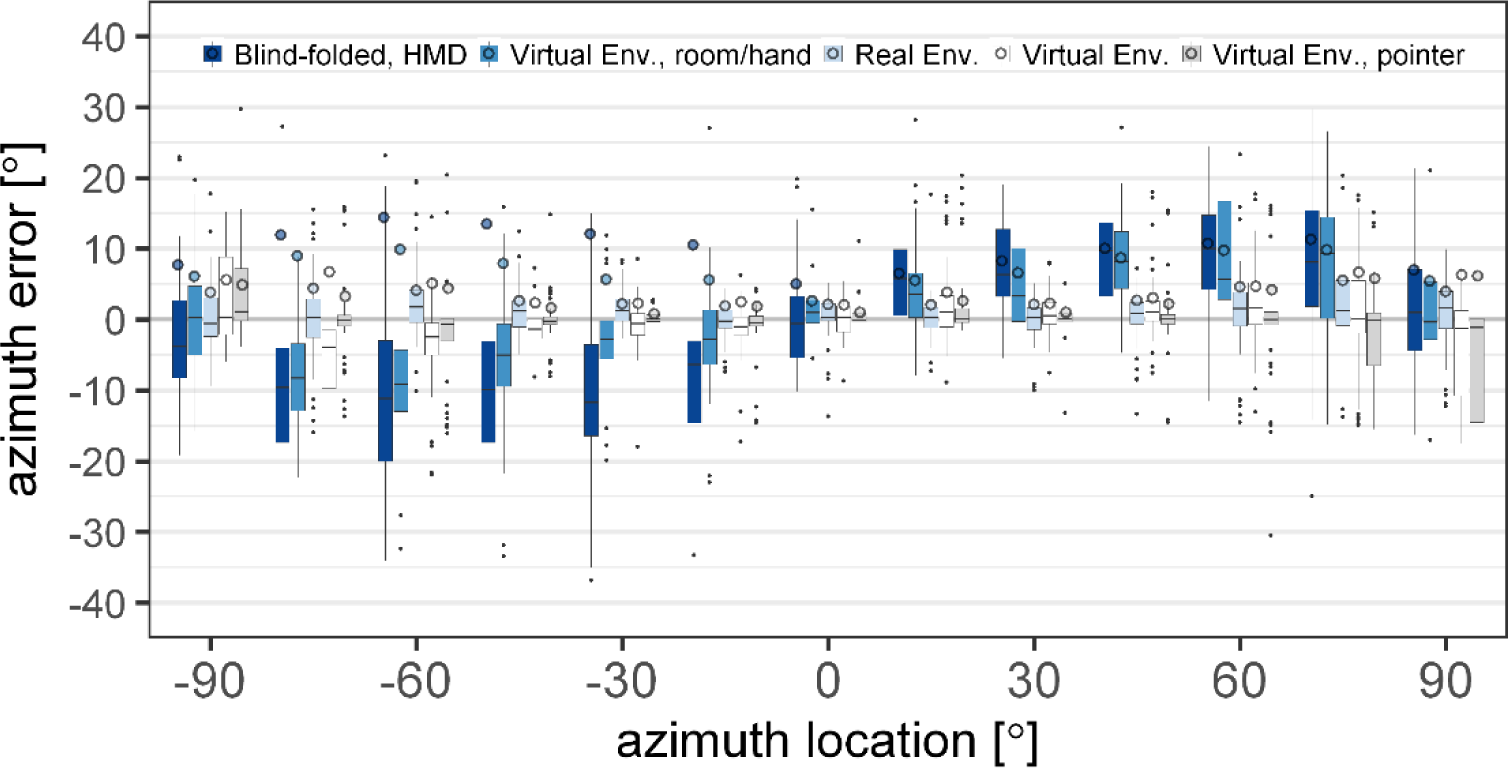
Mean absolute (circles) and signed (boxplots) azimuth error for acoustic localization with varying visual information in the virtual environment (VE) and the real environment (RE). The error is shown over the thirteen azimuth angles in the horizontal plane (0° elevation). The boxplots indicate the median (line) and the first and third quartile. The whiskers extend to 1.5 times the interquartile range.

The analysis of the linear mixed-effects model of the absolute azimuth error showed that both main effects of azimuth source location [F(12,3176)=23.17, p<0.0001] and conditions [F(4,3176)=223.88, p<0.0001] as well as their interaction [F(48,3176)=4.57, p<0.0001] had a significant effect on the azimuth error. A post-hoc analysis was performed to investigate the effect of the room and hand position information, the loudspeaker locations, and the laser pointer for aided pointing on the perceived sound source.

A significant decrease in error was found when comparing the blind-folded HMD condition (dark blue) with the condition where the subjects had visual information of room and hand position (blue) at azimuth location between −15° and −75° [p<0.039]. No significant change in error was found at −90° [t(3176)=1.61, p=1], at the right side of the subjects [p=1] nor for the frontal source [t(3176)=2.34, p=0.25].

When also visual information of the loudspeaker locations was provided (white), the subjects generally pointed towards the correct loudspeaker. When the loudspeaker locations were provided, the azimuth localization error decreased in comparison to the condition where only room and hand-location information were given. The reduction in error was significant for azimuth locations in the left hemifield between −15° and −60° [p<0.03] and in the right hemifield between 30° and 75° [p<0.02]. The lateral sources on the left [p>0.34] and on the right [p=1], as well as sources close to the midline at 0° and 15° [p=1] were not found significantly different in the two conditions with and without visual representations of the loudspeakers.

When the laser pointer was shown in the VE (grey), the absolute azimuth error was not found to be different from the condition without the laser pointer (white) [p=1], except for the azimuth angle −75° [t(3176)=3.42, p=0.0084]. Comparing the VE without the laser pointer (white) and the RE (light blue) showed no significant difference of the azimuth error at any of the source locations on the horizontal plane [p>0.28].

### Influence of HMD on elevation error

Figure 9 shows the error in elevation as a function of the elevation target location. The results for the blind-folded conditions are shown with the HMD (light grey) and without the HMD (dark blue). The analysis of the linear mixed-effects model including the two conditions, azimuth and elevation locations, revealed significant main effects of conditions [F(1,2049)=35.29, p<0.0001] and source elevations [F(2,2049)=8.28, p=0.0003] but no effect of the azimuth locations [F(6,2049)=1.98, p=0.0653]. The interactions of the source locations with the conditions were not found to be significant. The mean increase in elevation error between the conditions with and without the HMD was 1.8°.

**Fig 9.**
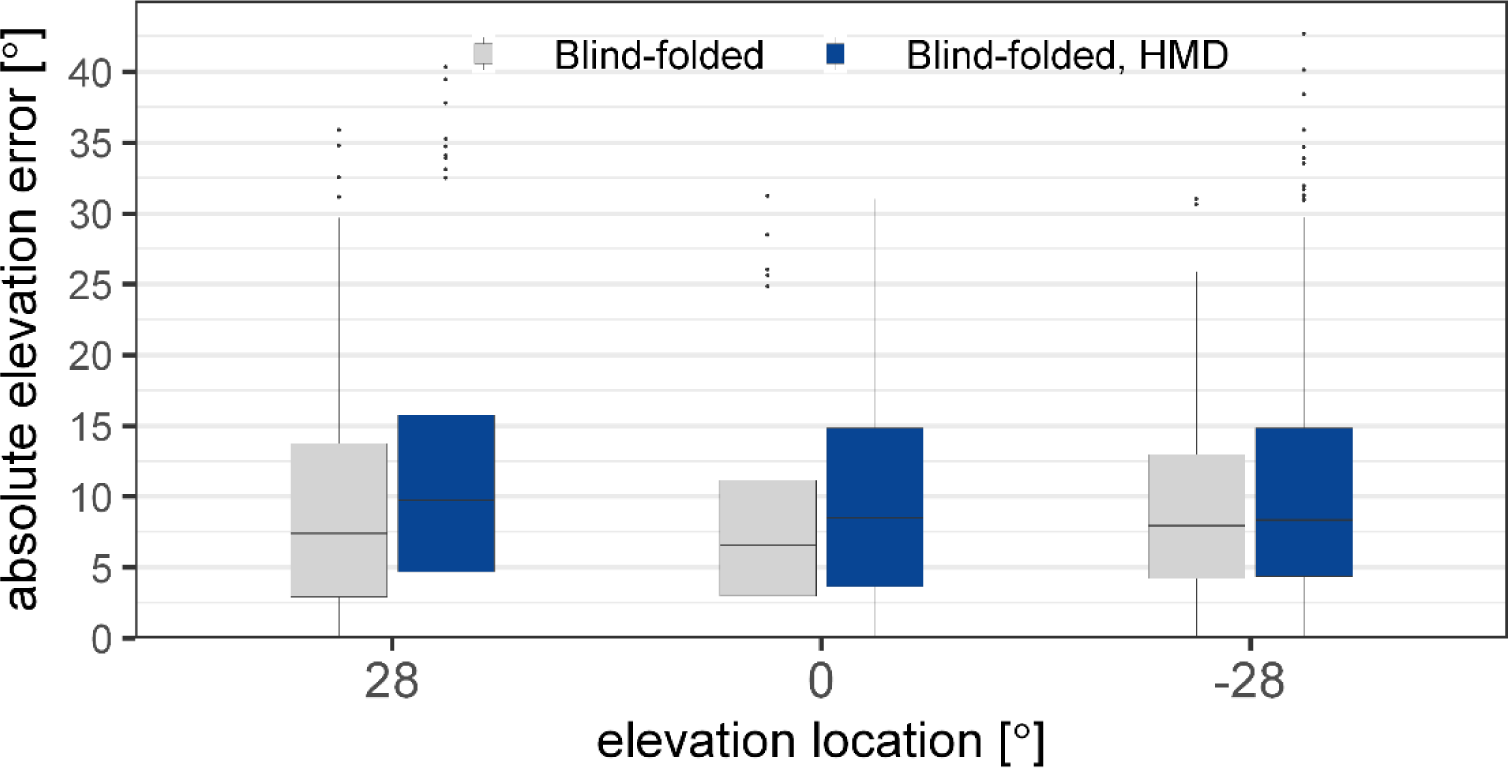
Absolute elevation error in degrees for acoustic localization for blind-folded subjects with (dark blue) and without (light grey) the head mounted display (HMD). The error is shown over the three elevation angles and includes the sources from all azimuth locations. The boxplots indicate the median (line) and the first and third quartile. The whiskers extend to 1.5 times the interquartile range.

### Influence of visual information on elevation error

Figure 10 shows the absolute elevation error at the three source elevations for the five conditions with varying visual information. The statistical analysis showed significant main effects of the conditions [F(4,5123)=289.44, p<0.0001] and the azimuth locations [F(6,5123)=9.47, p<0.0001], but no effect of the elevation locations [F(2,5123)=2.04, p=0.13]. The interaction between the conditions and the locations was found to be significant for the elevation [F(8,5123)=5.73, p<0.0001] and non-significant for the azimuth [F(24,5123)=1.43, p=0.0779].

**Fig 10.**
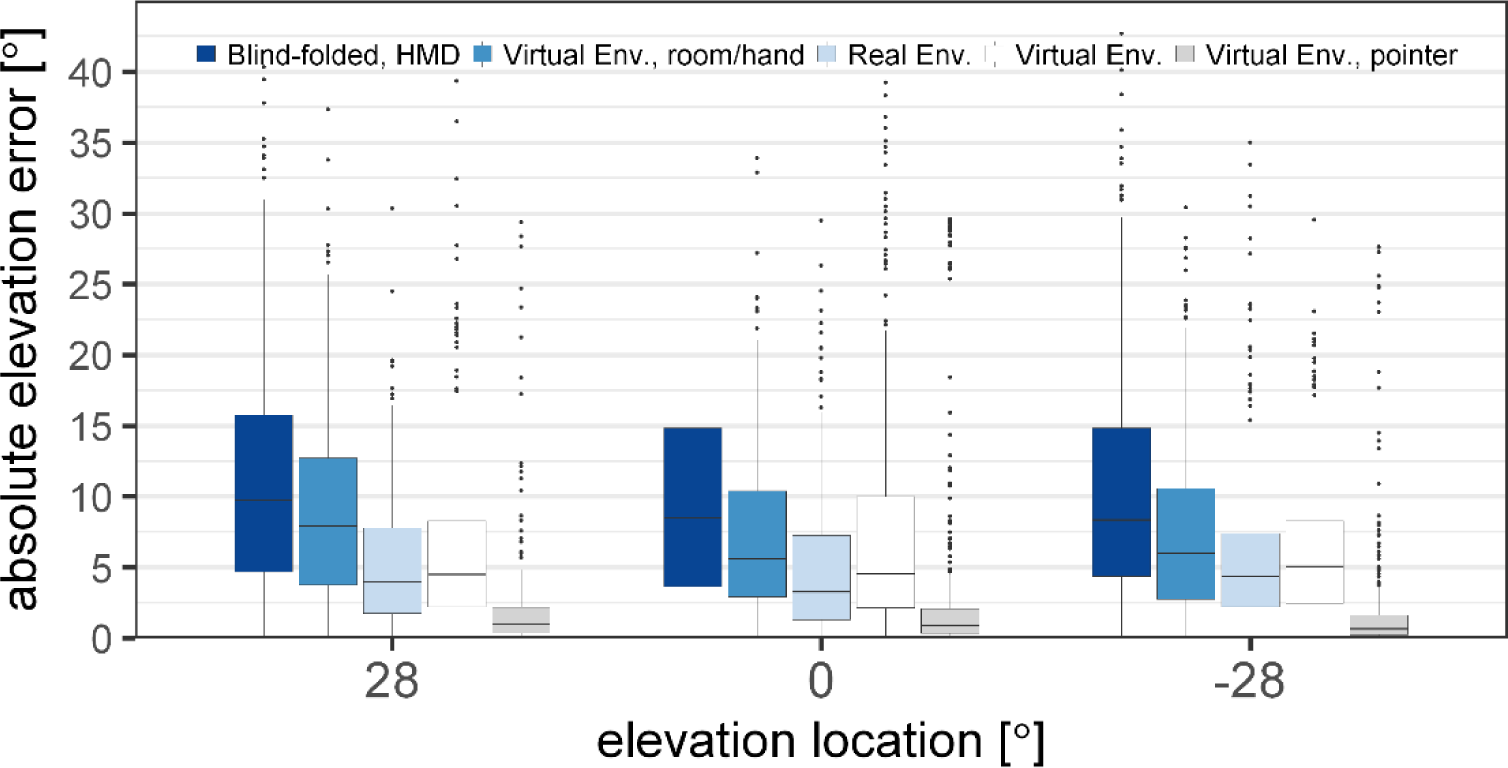
Absolute elevation error for acoustic localization with varying visual information in the virtual environment (VE) and the real environment (RE). The error is shown over the three elevation angles and includes the sources from all azimuth locations. The boxplots indicate the median (line) and the first and third quartile. The whiskers extend to 1.5 times the interquartile range.

The post-hoc analysis showed a significant drop in error between the blind-folded condition (dark blue) and the condition with visual room and hand-position information (blue) for all source elevations [p<0.0001]. The mean decrease of the error was 2.1° (from 10.2° to 8.1°). When the loudspeakers were visualized (white) in addition to the room and hand position (blue), the elevation error was found to further decrease significantly for the elevated sources [p<0.0018], but not for the sources in the horizontal plane [t(5123)=-0.18, p=1]. When the laser pointer was employed (grey), the absolute elevation error was lowest (2.4°) and significantly different from the error in the VE condition without the laser pointer (white) [p<0.0001]. The comparison of the elevation error in the RE (light blue) and the VE (white) revealed a significantly larger error for the sources at an elevation angle of 0° [t(5123)=-5.64, p<0.0001] but not for the sources above [t(5123)=-2.04, p=0.16] and below [t(5123)=-0.42, p=1] the horizontal plane.

## Discussion

### Degraded sound localization with HMD

To localize a sound in azimuth direction, the auditory system relies on the interaural disparities of time and level (ITD and ILD) [23]. The interaural disparities in the present study were found to be larger when measured with the HMD than without the HMD, consistent with results from the study of Gupta et al. [5]. Since Hafter & De Maio [24] found ITD just-noticeable differences (JNDs) between 20 and 40 µs for lateral sources, the differences in ITD of up to 60 µs induced by the HMD (see Figure 4) might be perceptible. Likewise, the ILD changes of up to 6 dB induced by the HMD (see Figure 4) were found to be above the JND of 2 dB [25, 26].

The effect of the HMD on the perceived location of acoustic stimuli in an anechoic environment was investigated in the blind-folded conditions. The error without the HMD was comparable in azimuth and elevation to values reported in earlier studies [e.g. 27]. No difference in azimuth localization error with and without the HMD was found for sources at or around the median plane, which is consistent with the small errors of ITDs and ILDs induced by the HMD. However, for lateral sources, the azimuth error was larger with the HMD than without the HMD, which is a consequence of the larger binaural disparities caused by the HMD. However, the increase in localization error was larger on the left side than on the right side. A comparable difference was also observed in the visual pointing experiment with larger errors on the left than on the right side when wearing the HMD (see Figure 6). Thus, there seem to be additional factors contributing to the localization error beyond the acoustical error induced by the HMD. Possibly, the HMD represents an obstacle when pointing with the right hand to the left hemisphere, resulting in a larger pointing error.

To localize sources in elevation, the auditory system mainly relies on direction dependent spectral cues provided by acoustical filtering of the pinnae, head and body [28]. Recent studies investigated the effect of HMDs on HRTFs in a similar way as in the current study and showed spectral perturbations up to about 6 dB at high frequencies [5, 6]. In the current study comparable spectral perturbations were found as well as an increase in elevation error by about 2°. However, it has been shown that spectral differences do not correlate well with elevation localization errors [29] but that elevation perception is based on multi-feature, template-based matching [30–32].

### Visual information influences sound localization in VR

In the virtual condition representing the simulated anechoic room without loudspeakers, a reference frame was provided by the room and the subjects could see the pointer. The sound localization error was found to be smaller with this visual information than in the blind-folded condition with the HMD. The contributions of the visual information about the hand position and about the room dimensions cannot be separated, since the current study was not designed to distinguish between these two visual aspects. Tabry et al. [10] also observed lower errors of both azimuth and elevation sound localization in conditions similar to those in the current study whereby real visual information of the subjects’ body and of the room were presented instead of virtual visual information. Thus, the amount of visual information provided in the current study may be considered to resemble those provided in a real environment. However, Tabry et al. [10] found substantially larger elevation errors than in the present study both in the condition with and without visual information. The smaller elevation errors found in the present study might be due to the limited set of elevation source locations (−28°, 0°, 28°) than in the study of Tabry et al. [10], where the range of elevations was between −37.5° and 67.5°. Subjects might be able to learn possible source locations which can improve the localization accuracy [33].

When the source locations were visible, the azimuth and elevation errors decreased by 3° and 1.5°, respectively, consistent with results obtained in real environments [34]. No improvement in localization accuracy was found in azimuth for frontal sources and in elevation for sources in the horizontal plane, because the auditory localization accuracy, even without visible source locations, was already high compared to those that are away from the midline and horizontal plane. However, there was likely a high visual bias towards the loudspeakers in this condition. Thus, mainly pointing accuracy and not sound localization accuracy was measured. The location priors have an even higher impact on the elevation than on the azimuth accuracy since only three elevation locations were used.

The additional information provided by the visual feedback from the laser pointer had only a negligible effect on the localization accuracy in azimuth, but a clear effect on the elevation accuracy. The elevation error was larger when no laser pointer was visible which might be partly due to the shape of the VR controller. The shape of the controller led to a biased pointing direction. The bias correction as described above (see Figure 2) was intended to reduce the influence of the controller. However, even though the subjects were asked to always hold the controller in the same way, the controller positioning might have varied leading to a larger pointing error when no visual feedback of the laser pointer was provided. Even though the effect of the laser pointer on the mean azimuth error was negligible, the variance of the subjects’ responses decreased when the visual feedback of the pointing direction was available. Thus, the responses with the laser pointer become more reliable.

Comparing the localization error in the real and the virtual environments showed no differences in terms of the azimuth error. The elevation error was significantly increased at 0° elevation and was found to be slightly, but not significantly, larger above the horizontal plane even though not significantly. Below the horizontal plane no difference between RE and the VE was found. Even though the provided visual information was supposed to be the same in the two environments, some differences were unavoidable. In the real loudspeaker environment the subjects could see their arms, but not in the VE, where only the controller was visible. However, Van Beers et al. [35] showed that the visual feedback of the arm does not seem to increase visual object localization accuracy compared to the situation when only the finger is visible. Nevertheless, for pointing to sources on and above the horizontal plane the arm might have been helpful visual information for more accurate pointing, however, no such evidence was found in the visual pointing experiment.

### Potential training effects

The subjects did not receive training before starting the experiment, but were introduced to the task and the controller. Previous studies indicated that training can improve sound localization accuracy [16,36,37]. Since the subjects could not be introduced to the visual loudspeaker environment beforehand, no training was provided to avoid potential benefits in certain conditions but not in others. It is possible that the localization performance of the subjects improved throughout the course of the experiment. To minimize this effect, the conditions were presented in a random order within the experimental blocks. The random-order presentation avoided a bias within the blocks but could not exclude an inter-block bias. Such inter-block bias was unavoidable because visual information needed to be revealed to the subjects reflecting an increase of information content.

### Implications for VR in hearing research

VR glasses in combination with audio reproduction techniques may allow for novel ways of conducting research in simulated realistic environments. While previous research typically involved simple audio-visual information, with VR, research can be conducted in ecologically more valid environments while maintaining controllability and reproducibility of the experiment. Even though the localization error increased with the HMD in the blind-folded condition, these errors may not be noticeable in realistic environments including visual information. This might also be the case for hearing-impaired subjects for whom sound localization accuracy is commonly reduced compared to normal-hearing subjects [38].

Even though only a single device, the HTC Vive, was investigated in the current study the findings may generalize with regards to other commercial virtual reality glasses. It has been shown that other HMDs, as for example the Oculus Rift (Oculus VR LLC, Menlo Park, CA) or the Microsoft HoloLens (Microsoft, Redmond, WA), lead to comparable or even smaller acoustic perturbations [5, 6]. Thus, the sound localization error due to the HMD is likely to be comparable or lower than that of the HTC Vive. Visual reproduction and tracking specifications seem comparable between current commercial products.

## Conclusions

VR systems and loudspeaker-based audio reproduction allow for full immersion into an audio-visual scene. In the present study, sound source localization accuracy with an HMD providing a varying amount of visual information was investigated. A calibration system to align the real world and the virtual world was developed. Hand-pointing accuracy to visual targets was evaluated using commercially available controllers. The accuracy of the hand pointing to visual targets was found to be high in the azimuth direction, whereas a large bias was found in terms of elevation accuracy due to the shape of the controller. The sound localization experiment demonstrated a small detrimental effect of the HMD on the sound localization accuracy. While the azimuth error induced by wearing the HMD was negligible in the frontal area, it was significant at more lateral sound source locations which correlates with changes in binaural disparities. However, the error was found to be larger on the left than on the right side in both the acoustic and the visual pointing experiment. Thus, the error was not purely of acoustic nature but also due to the HMD influencing the motion behavior. The elevation error was about 2° larger with the HMD for all azimuth and elevation directions.

Generally, the sound localization accuracy was found to be higher when visual information was available than in the conditions without visual information. The lowest accuracy was found when the subjects were blind-folded and a significant improvement was found for both azimuth and elevation when room and hand position information were provided. An additional laser pointer for pointing guidance did not lead to an improvement of azimuth localization but an improved elevation localization.

